# Diagnostic value of chest ultrasound in children with cystic fibrosis

**DOI:** 10.1101/604942

**Authors:** Lidia Strzelczuk-Judka, Irena Wojsyk-Banaszak, Aleksandra Zakrzewska, Katarzyna Jonczyk-Potoczna

## Abstract

Cystic fibrosis (CF) is one of the most common genetic disorders in the Caucasian population. The disease has a progressive course and leads to reduced life quality and life expectancy. Standard diagnostic procedures used in the monitoring of CF patients, include methods exposing patients to the ionizing radiation. With increasing life expectancy in CF the cumulative dose of ionising radiation increases, prompting clinicians’ search for safer imaging studies. Despite its safety and availability lung ultrasound (LUS) is not routinely used in the diagnostic evaluation of CF patients.

The aim of the study was to evaluate the diagnostic value of LUS in children with CF compared to chest X-ray, and to assess the diagnostic value of the recently developed LUS score - CF-USS (Cystic Fibrosis Ultrasound Score).

LUS was performed in 48 CF children aged from 5 to 18 years (24 girls and 24 boys). LUS consisted in the assessment of the pleura, lung sliding, A-line and B-line artifacts, “lung rockets”, alveolar consolidations, air bronchogram and pleural effusion. Chest radiography was performed in all patients and analyzed according to the modified Chrispin-Norman score. LUS was analyzed according to CF-USS.

Correlation between the CF-USS and the modified Chrispin-Norman scores were moderate (R=0.52, p=0.0002) and strong in control studies. In 75% of patients undergoing LUS, small areas of subpleural consolidations were observed, not visible on X-rays. At the same time, LUS was not sensitive enough to visualize bronchial pathology, which plays an important role in assessing the disease progression.

**Conclusions:** LUS constitutes an invaluable tool for the diagnosis of subpleural consolidations. CF-USS results correlate with conventional x-ray modified Chrispin–Norman score. LUS should be considered an accessory radiographic examination in the monitoring of CF patients, and CF-USS may provide clinicians with valuable information concerning the disease progression.

## Introduction

Cystic fibrosis (CF) is one of the most common autosomal recessive hereditary life-shortening disorders in Caucasian populations [1,2]. The disease is caused by the mutation of gene coding CFTR protein (*Cystic Fibrosis Transmembrane Conductance Regulator*), leading to the production of dense mucus in the airways and exocrine glands and the impairment of their function. The main affected systems comprise respiratory and digestive systems, and the chronic pulmonary disease remains the main cause of morbidity and one of the most important prognostic factors in CF [1,3,6]. Chronic inflammation due to impaired mucocilliary clearance and mucus impaction in the airways results in bronchiectasis and progressive lung tissue destruction [5].

Lung evaluation in CF patients traditionally avails of imaging studies and among these the most commonly used remains chest x-ray. Early in the course of disease the radiologic picture might reveal no abnormalities. Along with the disease progression lung hyperinflation and increased bronchial markings appear, followed by chest infiltrates, atelectasis and bronchiectasis [1,6].

The need for objective tools for the evaluation of patients has prompted the development of x-ray scoring systems including Brasfield score [7] Northern score [4] Chrispin-Norman score [8] and its modified version [9]. These scoring systems are used for the monitoring of disease progression, evaluation of different therapies as well comparison of patients’ outcomes between the treatment centres [4,8-14].

The most accurate radiographic diagnostic modality in CF, so called „golden standard” that allows for qualitative and quantitative evaluation of lung involvement, even very early in the course of the disease remains computed tomography (CT) [6]. CT due to its high resolution allows visualisation of the detailes that are not visible in the plain chest x-ray [10]. In CF patients CT enables visualisation of bronchial wall and peribronchial thickening, intralobular nodules, bronchiolitis, so called „tree in bud” sign, air trapping, bronchiectasis, mucus impaction, microabscesses, infiltrates, atelectasis, enlarged lymph nodes and widening of pulmonary artery with narrowing of peripheral vessels [5,15,16]. The role of CT in CF patients was confirmed in studies reporting on correlation of CT scans with patients outcomes [17]. For quantitative, objective evaluation of CT results in CF patients scoring systems were also developed with the most popular Bhalla score [18].

Disadvantage of CT scanning is a relatively high dose of ionising radiation. The risks of cancer related to lifetime exposure to radiation made clinicians look for imaging modalities with the lowest or ideally no radiation [19]. Ultrasound (US) is currently one of the most important and most frequently used imaging techniques [20]. Considering this, lung ultrasound (LUS) as a safe, non-invasive, widely available and cheap technique might constitute an important tool in the diagnostic protocols of children with CF [21]. Despite this fact there are few existing reports on LUS application in CF patients. There are only two reports published as abstracts by Ciuca et al on LUS in CF as compared to CT scans [22,23].

LUS examination comprise evaluation of pleural line and lung sliding [24-28], analysis of the artefacts that are present in normal lung, like „the bat sign” [24,25,29,30] and the A-line artefacts [25,28,31,32] as well as in pathological conditions (the B-line, Z-line and I-line artefacts) and evaluation of thoracic wall structures. The B-lines are vertical, well defined hyperechogenic lines, arising from the pleural line, spreading out without fading to the edge of the screen, similar to laser beam or „comet tail” artefact [25,33,34]. Multiple B-lines are typical for interstitial lung disease [35-37]. Seen together they are described as „lung rockets” artefact [28,32,37]. Multiple coalescent B-lines in the absence of A-lines with visible lung sliding constitute so called „white lung” image [37-39]. Alveolar consolidation can be diagnosed with LUS provided their peripheral localisation and according to the literature reports that is the case in up to 98.5% of cases [24,29,32].

## Objective

The aim of our report was to evaluate the diagnostic value of chest ultrasound in children with CF as compared to plain x-ray, as well as to assess the diagnostic value of the recently developed LUS score - CF-USS (Cystic Fibrosis Ultrasound Score).

## Material

We enrolled 48 Caucasian patients (24 males) aged 5 to 18 years diagnosed with CF who were admitted to the Pulmonology Department for scheduled annual diagnostic workup. Patients underwent chest ultrasound and plain x-ray, and the time interval between the studies were not longer than 72 hours.

In all the studied children CF was confirmed by two positive sweat test results ant genetic studies (two pathogenic mutations). All the patients and their parents gave informed consent for the study. Exclusion criteria comprised severe immunosuppression, lack of consent and time interval longer than 72 hours between the studies. The study design was accepted by the Bioethical Committee of Poznań University of Medical Sciences. In Table 1 we presented the characteristic of the studied patients.

**Table 1.** Patients characteristic

|  |  |  |
| --- | --- | --- |
| Number of patients | 48 |  |
| Males/females | 24 / 24 |  |
| | Mean ± SD | Median (minimum-maximum) |
| Age (years) | 11.9 ± 3.9 | 12 (5-18) |
| BMI | 16.7 ± 2.9 | 15.7 (10.2-24.2) |
| FEV<sub>1</sub> (L) | 1.9 ± 0.8 | 1.7 (0.4 - 4.3) |
| BMI <3c N(%) | 7 (14.6) |  |
| Pancreatic sufficient N (%) | 9 (18.8) |  |
| F508del homozygous N(%) | 21 (43.8) |  |
| F508del heterozygous N(%) | 18 (37.5) |  |
| Chronically infected with *P. aeruginosa* | 21 (43.8) |  |
| FEV<sub>1</sub> ≥ 80% normal value | 32 (66.7) | |
| FEV<sub>1</sub> ≤ 40% normal value | 1 (2.1) | |

## Methods

### Radiographic imaging

X-rays were performed with an analogue apparatus Axiom Iconos R 100 (Siemens Healthcare), in posteroanterior projection during suspended inspiration. Technical parameters of the images (including use of grid, the source image receptor distance, dose of radiation) were individually adjusted for every studied patient in concordance with ALARA (As Low As Reasonably Achievable) principle in order to achieve best possible images using the lowest radiation dose. No lateral x-rays were performed. X-rays were independently evaluated by two board certified paediatric radiologists with experience in CF. Chest x-rays were evaluated using modified Chrispin-Norman score [9].

### Chest ultrasound (LUS)

Chest ultrasound was performed with iU22 apparatus (Philips, Biothel United States) using linear probe of 5 – 12 mHz (L12-5) frequency and depending on the patients’ age with either convex probe of 1 - 5 mHz (C5-1) frequency, convex probe of 4 - 9 mHz (C9-4) frequency or microconvex probe of 5 – 8 mHz (C8-5) frequency through longitudinal and transverse sections of anterior, lateral and posterior wall of the chest. Preliminary preset was soft tissue excluding artefact reduction options (SonoCT, XRes). Doppler imaging was used for the evaluation of vascularisation of the inflammatory changes.

Patients were examined in the sitting position. The studies were performed by two board certified paediatric radiologists with experience in LUS and CF.

In every patient we evaluated the quality (free flowing or organised, localization) and quantity (fluid layer in millimetres) of any fluid present in the pleural space, the shape and thickness of the pleural line, the lung sliding sign, A-lines and B-lines artefacts (their number, localisation and morphology, including single ones as well as „lung rockets” complexes and “white lung” images) and alveolar consolidations (their number, dimensions, localisation, morphology, presence of bronchogram and its characteristic (air or fluid) and vascularisation).

LUS results were classified according scoring system developed by the authors: CF-USS *(Cystic Fibrosis Ultrasound Score)* devised on the basis of modified Chrispin-Norman score and bronchiolitis score reported by Caiulo and collaborators [39,40]. Scores are calculated separately for anterior and posterior surface of right and left half of the thorax. Each part can be scored from 0 to 2 points for irregularities of pleural line, single and complex B-line artefacts, alveolar consolidations and the presence of fluid in the pleural space with the maximum score of ten for each part and 40 in total. The higher the score, the more advanced the disease process (Table 2).

**Table 2.** Cystic Fibrosis Ultrasound Score

| Characteristic | Intensity |  |  |
| --- | --- | --- | --- |
| **Pleural irregularities** | Absent | Present | Present+ pleural thickening |
| Right lung: anterior surface | 0 | 1 | 2 |
| posterior surface | 0 | 1 | 2 |
| Left lung: anterior surface | 0 | 1 | 2 |
| posterior surface | 0 | 1 | 2 |
| **Focal B-line artefacts** | Absent / few ($\leq 6$) | Some (7-14) | Many ($\geq 15$) |
| Right lung: anterior surface | 0 | 1 | 2 |
| posterior surface | 0 | 1 | 2 |
| Left lung: anterior surface | 0 | 1 | 2 |
| posterior surface | 0 | 1 | 2 |
| **Coalescent B-line artefacts** | Absent | Fused | „Lung rockets” |
| Right lung: anterior surface | 0 | 1 | 2 |
| posterior surface | 0 | 1 | 2 |
| Left lung: anterior surface | 0 | 1 | 2 |
| posterior surface | 0 | 1 | 2 |
| **Subpleural consolidations** | Absent ($\leq 6$) | Some (7-14) | Multiple or extensive ($\geq 15$) |
| Right lung: anterior surface | 0 | 1 | 2 |
| posterior surface | 0 | 1 | 2 |
| Left lung: anterior surface | 0 | 1 | 2 |
| posterior surface | 0 | 1 | 2 |
| **Pleural fluid** | Absent | Obliterating phreno – costal angle | In phreno – costal angle and along the chest wall |
| Right lung: anterior surface | 0 | 1 | 2 |
| posterior surface | 0 | 1 | 2 |
| Left lung: anterior surface | 0 | 1 | 2 |
| posterior surface | 0 | 1 | 2 |

### Statistical analysis

Statistical analysis was performed with Statistica software (version 12; StatSoft). Data distribution was evaluated with Shapiro-Wilk test. For data with normal distribution we used t Student test for paired and independent variables. For data that do not meet the normal distribution assumptions Spearman’s rank correlation coefficient was calculated. P value of <0.05 were considered statistically significant.

## Results

### Comparison of LUS and chest x-ray images

Pulmonary disease was evaluated radiologically with modified Chrispin – Norman score for x-rays and CF-USS score. The patients’ results are shown in Table 3, Figures 1 and 2. Statistical analysis has shown positive correlation between the two scoring systems (R Spearman= 0.52, p=0.0002) (Fig 3).

**Table 3.** Comparison of two scoring systems

| Parametr | Modified Chrispin – Norman score | CF-USS |
| --- | --- | --- |
| Number of patients | 48 | 48 |
| Mean ± SD | 8.7 ± 8.0 | 4.4 ± 4.1 |
| Median (minimum-maximum) | 7 (0-28) | 4 (0-16) |

**Fig 1.**
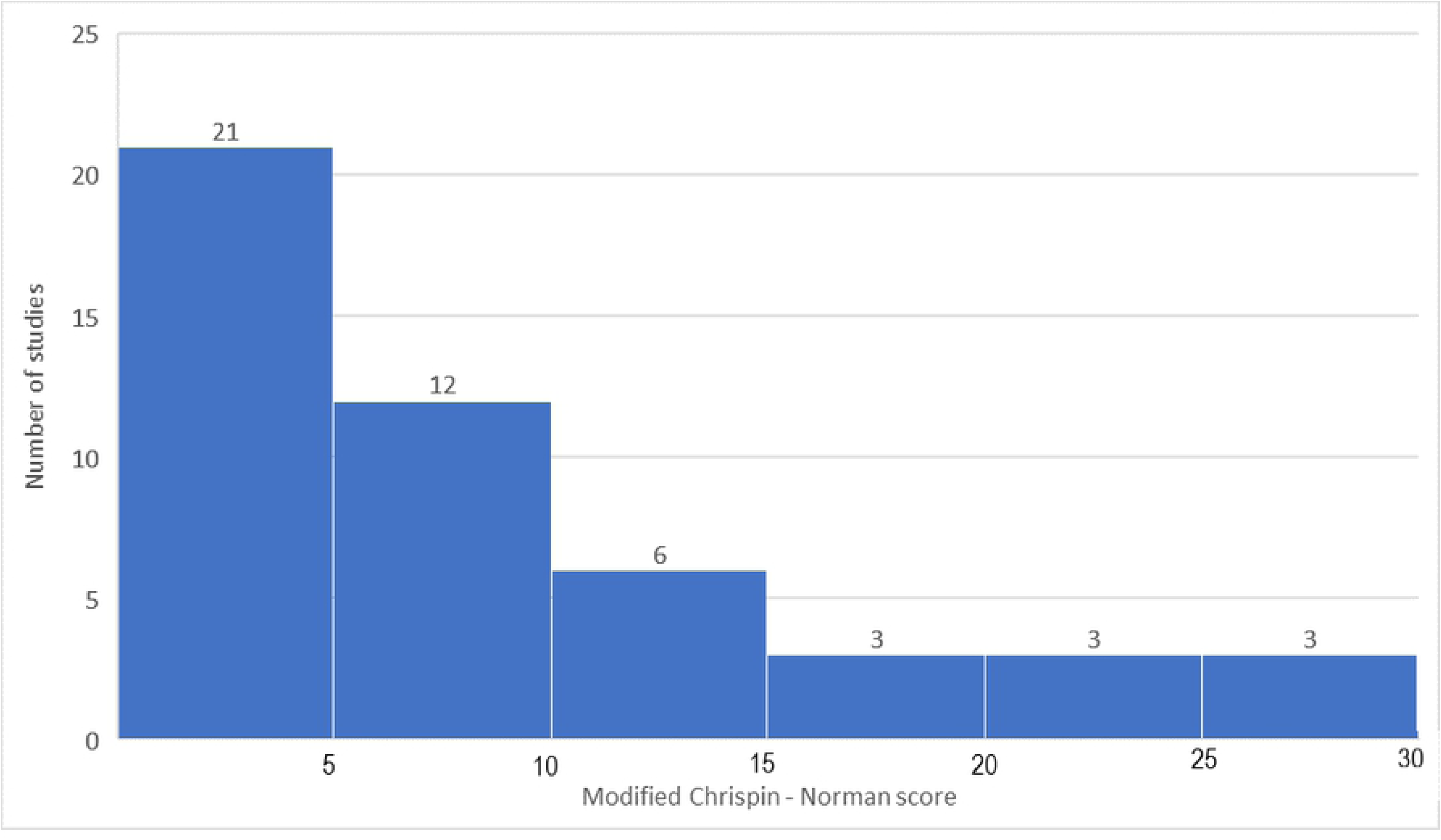
The modified Chrispin-Norman scores.

**Fig 2.**
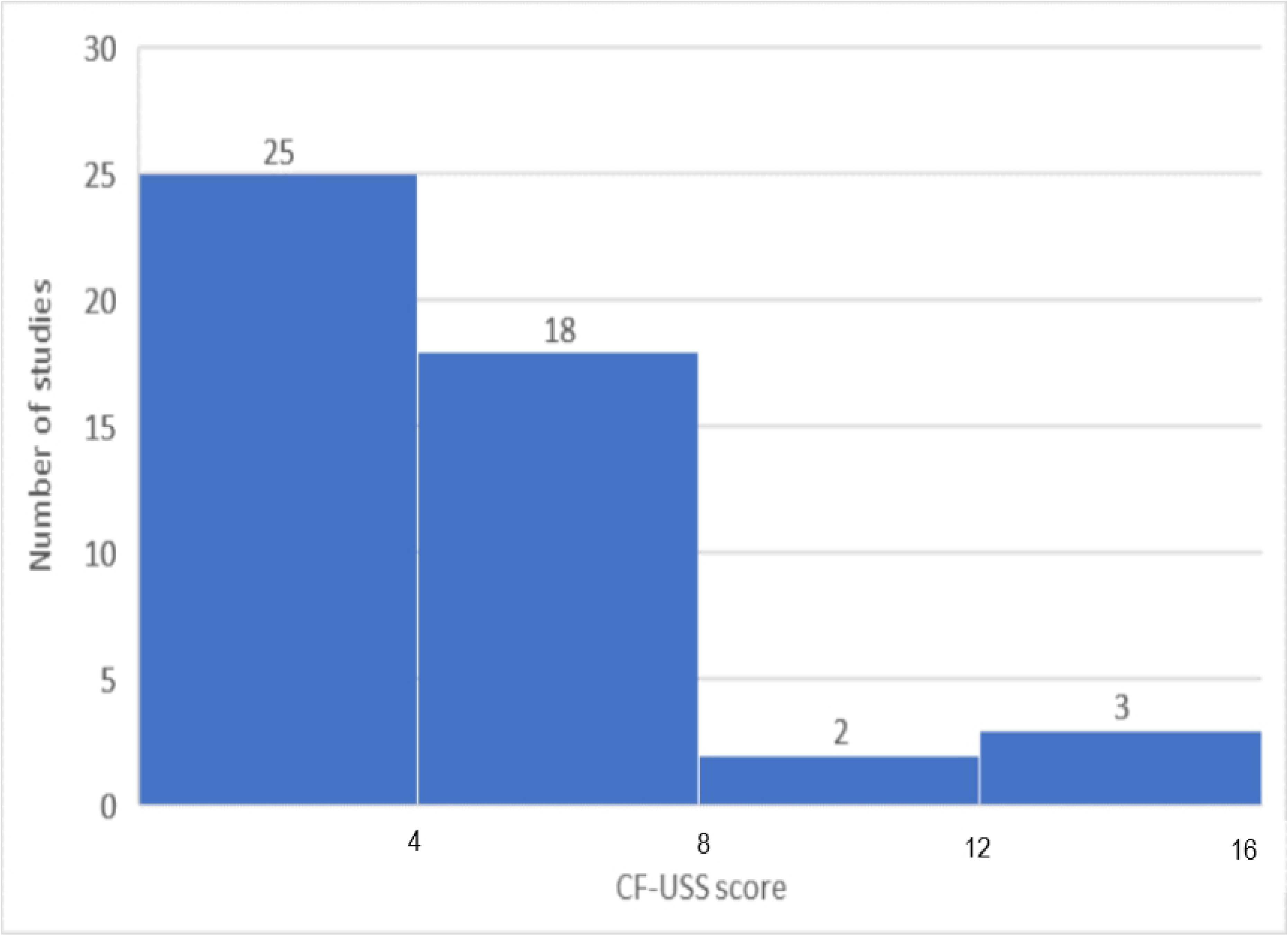
The CF-USS scores.

**Fig 3.**
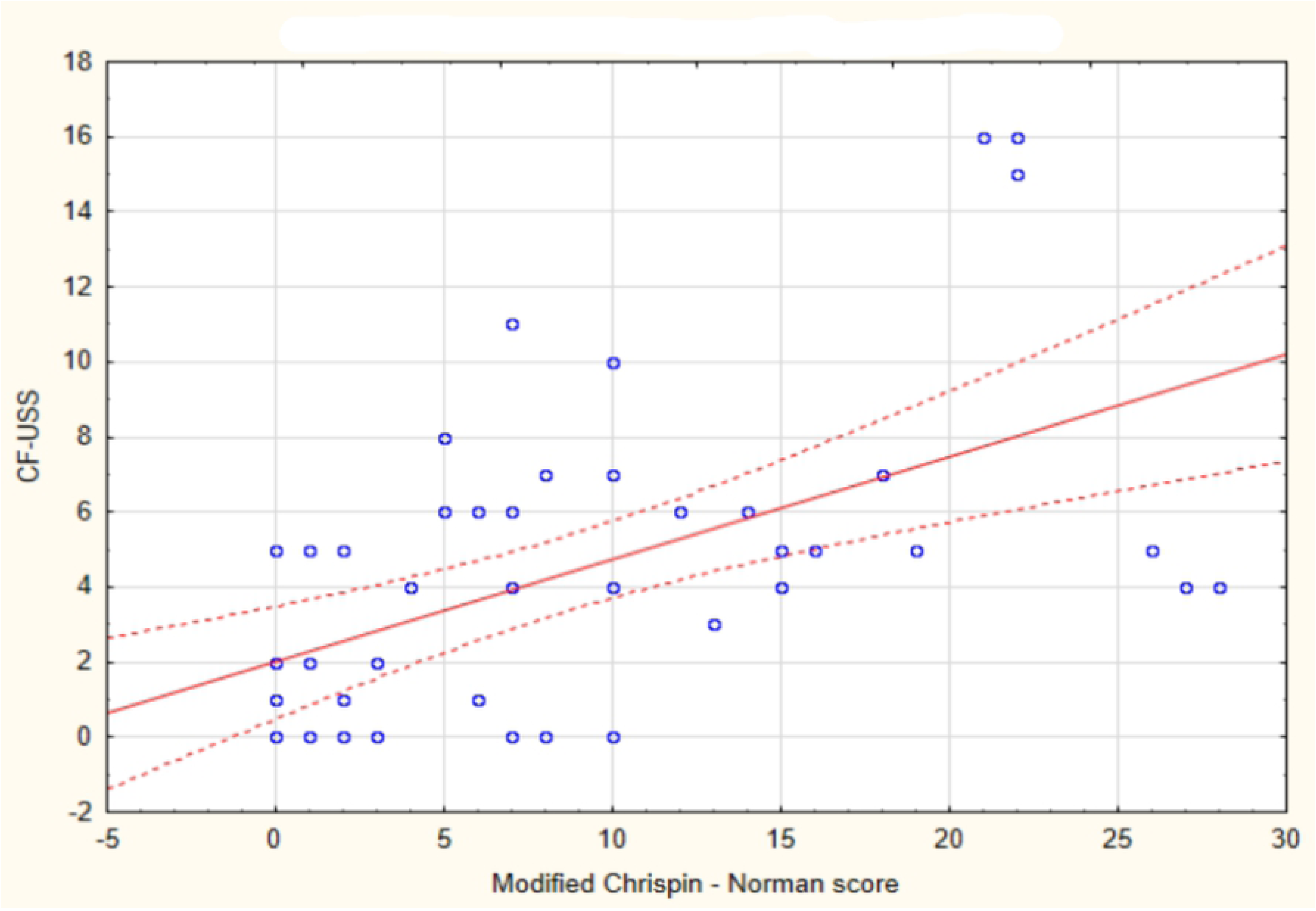
Correlation between CF-USS and the modified Chrispin-Norman scores.

Fine subpleural consolidation were seen in LUS in 36 patients (75%). Abnormalities seen in LUS in the studied patients classified according to CF-USS are presented in Table 4.

**Table 4.** Number of patients in the studied group with abnormalities according to the CF-USS score.

| Characteristic | Number of patients | Percentage (%) |
| --- | --- | --- |
| **Pleural irregularities** | 1 | 2 |
| **Focal B-line artefacts** |  |  |
| Some | 33 | 69 |
| Many | 4 | 8 |
| **Fused B-line artefacts** |  |  |
| Coalescent | 19 | 40 |
| „Lung rockets” | 1 | 2 |
| **Subpleural consolidations** |  |  |
| Few | 28 | 58 |
| Multiple or extensive | 8 | 17 |
| **Pleural fluid** |  |  |
| In costo – phrenical angle | 9 | 19 |
| And along the chest wall | 2 | 4 |

In nine patients LUS and x-ray were performed twice on two different occasions. There were no statistically significant differences between the x-rays in modified Chrispin – Norman score (15.22±2.71 vs. 10.78±3.00; p=0.06) and LUS in CF-USS score (7.56±1.58 vs. 5.33±1.86; p=0.29) (Fig 4 and Fig 5). Statistical analysis for the repeated LUS and x-ray examinations showed positive correlation for the two studies (R Spearman= 0.81, p=0.01) (Fig 6).

**Fig 4.**
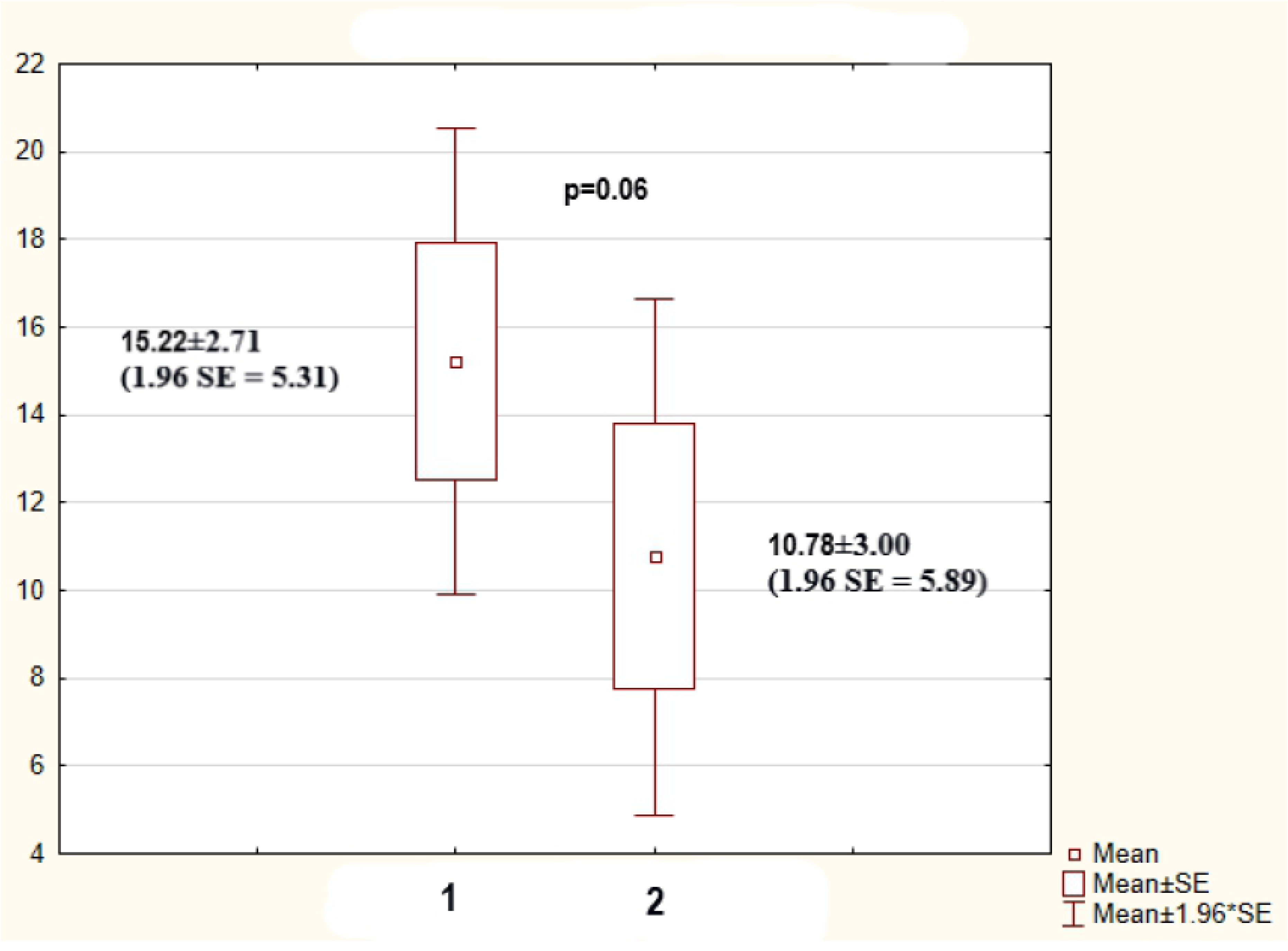
Modified Chrispin – Norman scores for the repeated studies. 1. Modified Chrispin – Norman scores in the first study; 2. Modified Chrispin – Norman scores in the repeated study

**Fig 5.**
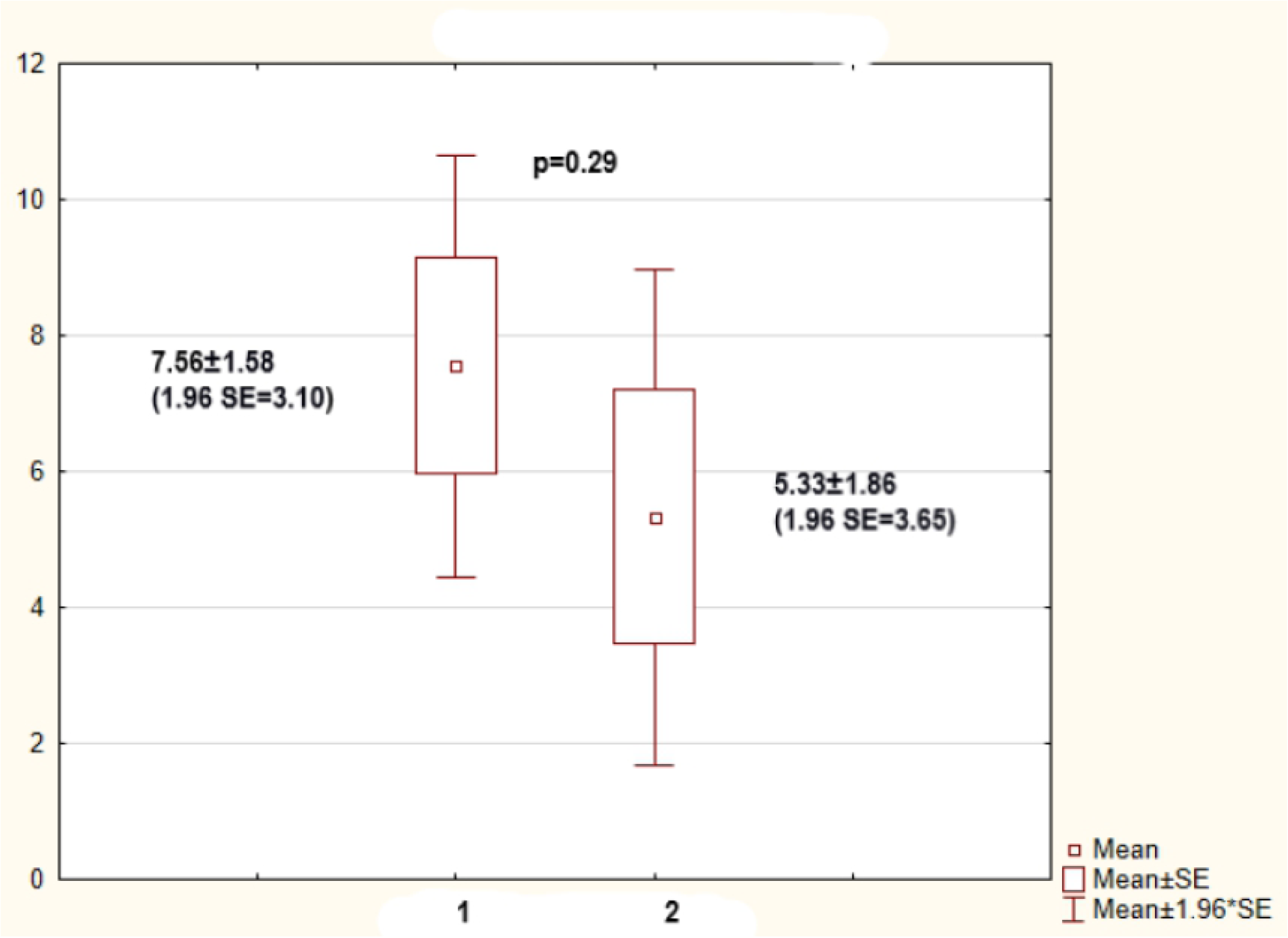
CF-USS scores for the repeated studies. 1. CF-USS scores in the first study; 2. CF-USS scores in the repeated study

**Fig 6.**
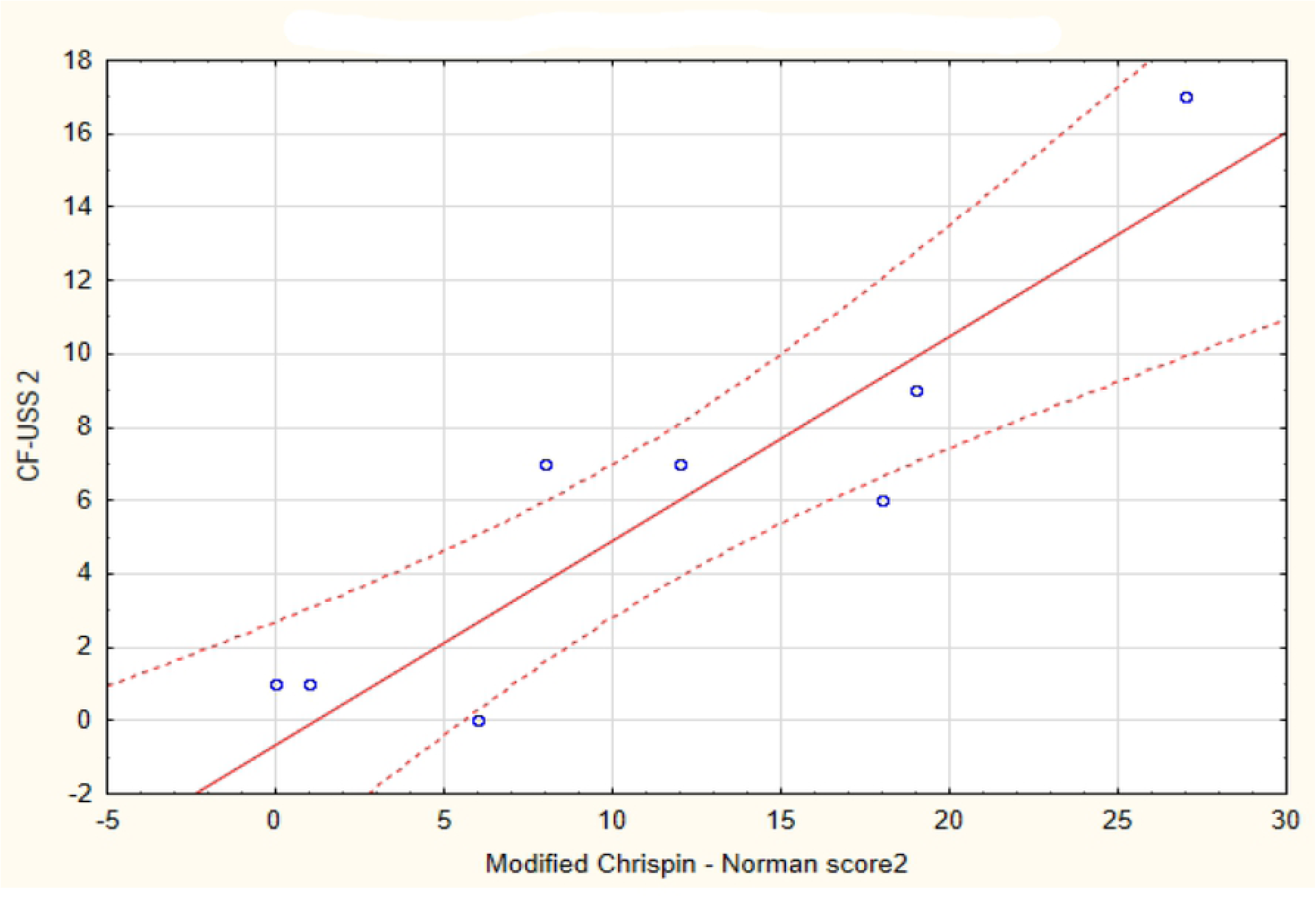
Correlation between the CF-USS score and the modified Chrispin-Norman score in the repeated studies. In the figures 7-11chest x-ray and LUS images of the studied patients are presented.

**Fig 7.**
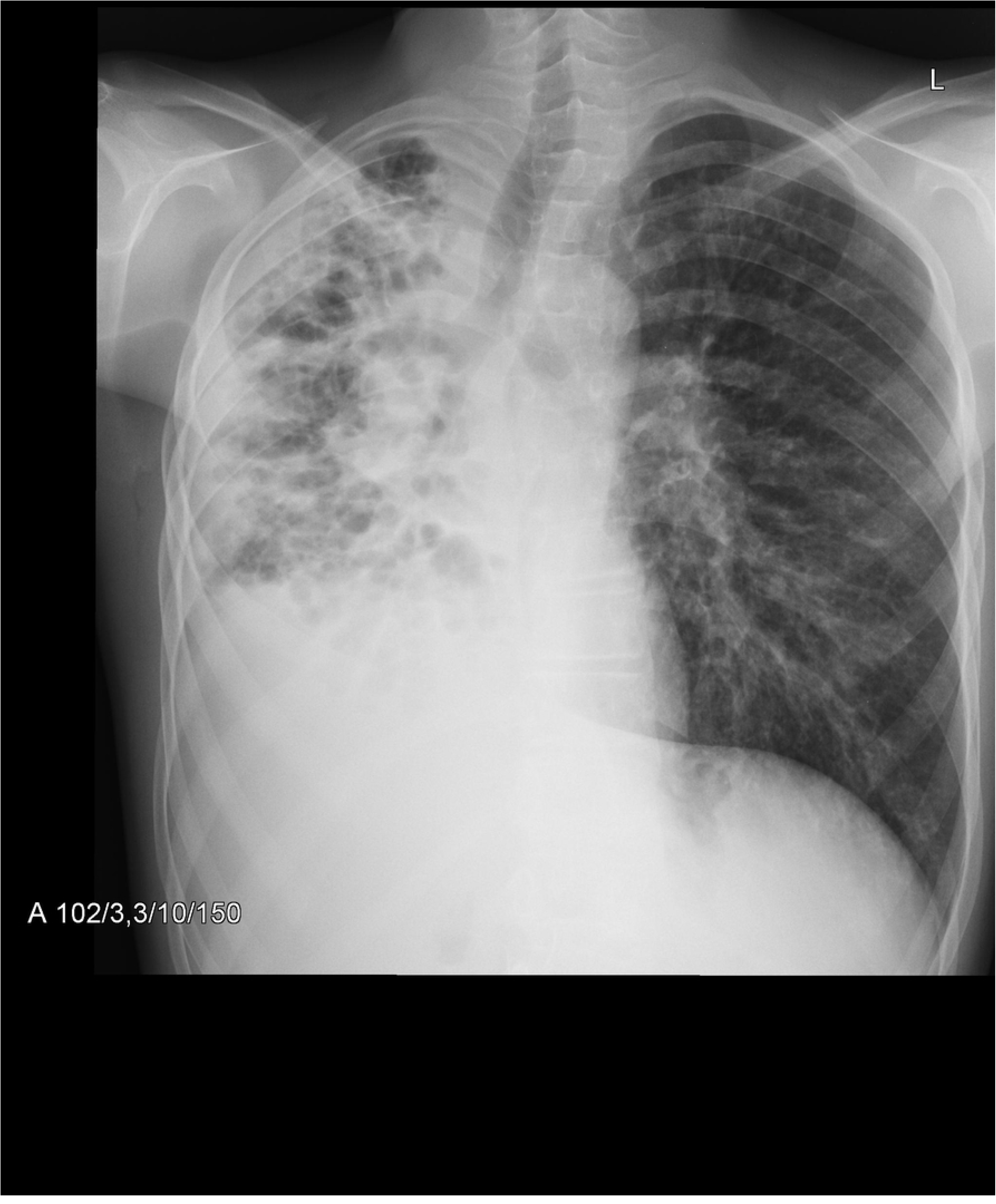
Mediastinal shift, atelectasis and pleural effusion on the right, linear and cystic opacities, bronchiectasis, consolidations. Hyperinflation of the left lung, with nodular and linear interstitial opacities. 27 points in modified Chrispin-Norman score.

**Fig 8.**
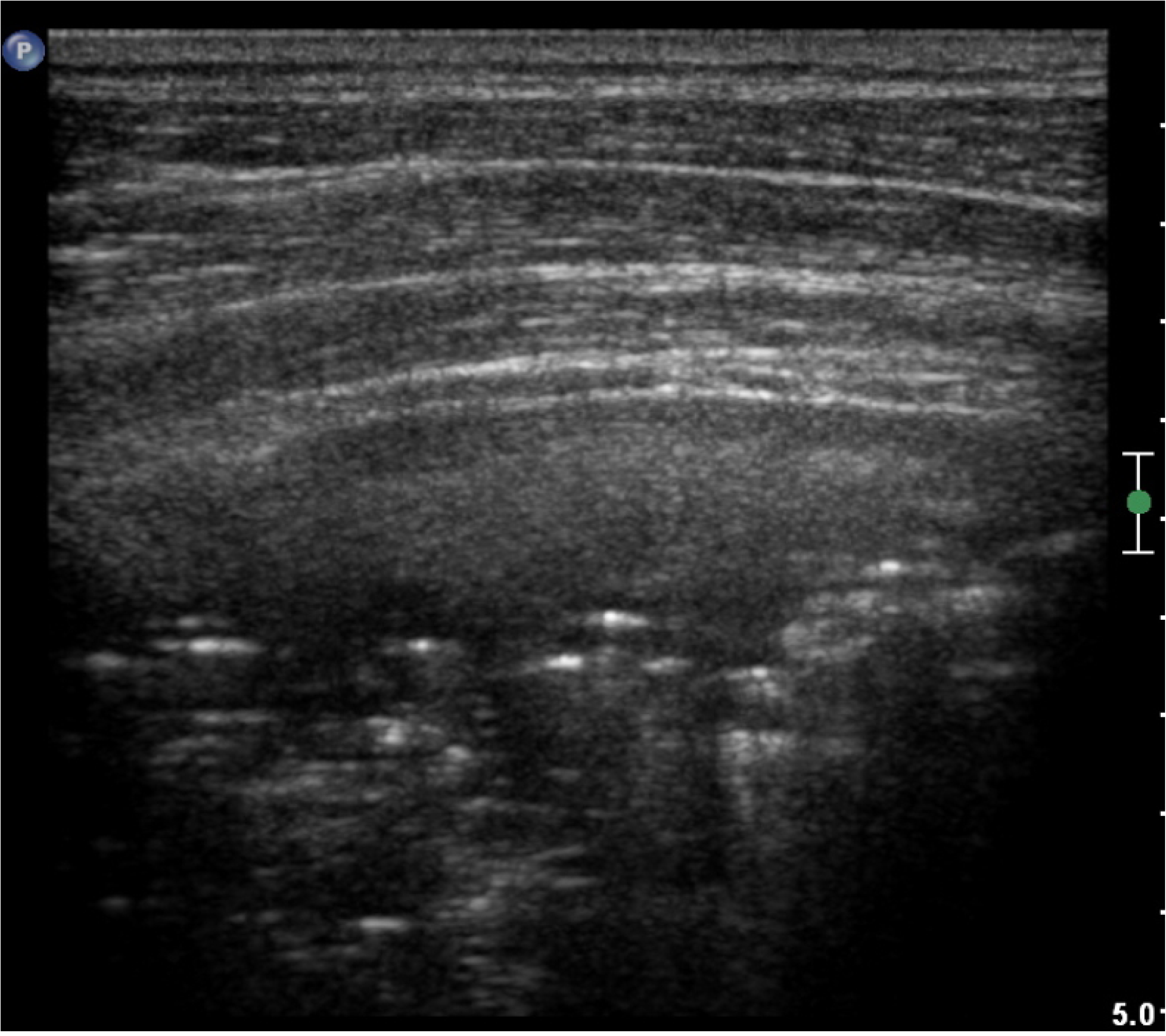
LUS image of the patient from Fig 7, linear probe. Regions of consolidation and atelectasis. 17 points in CF-USS.

**Fig 9.**
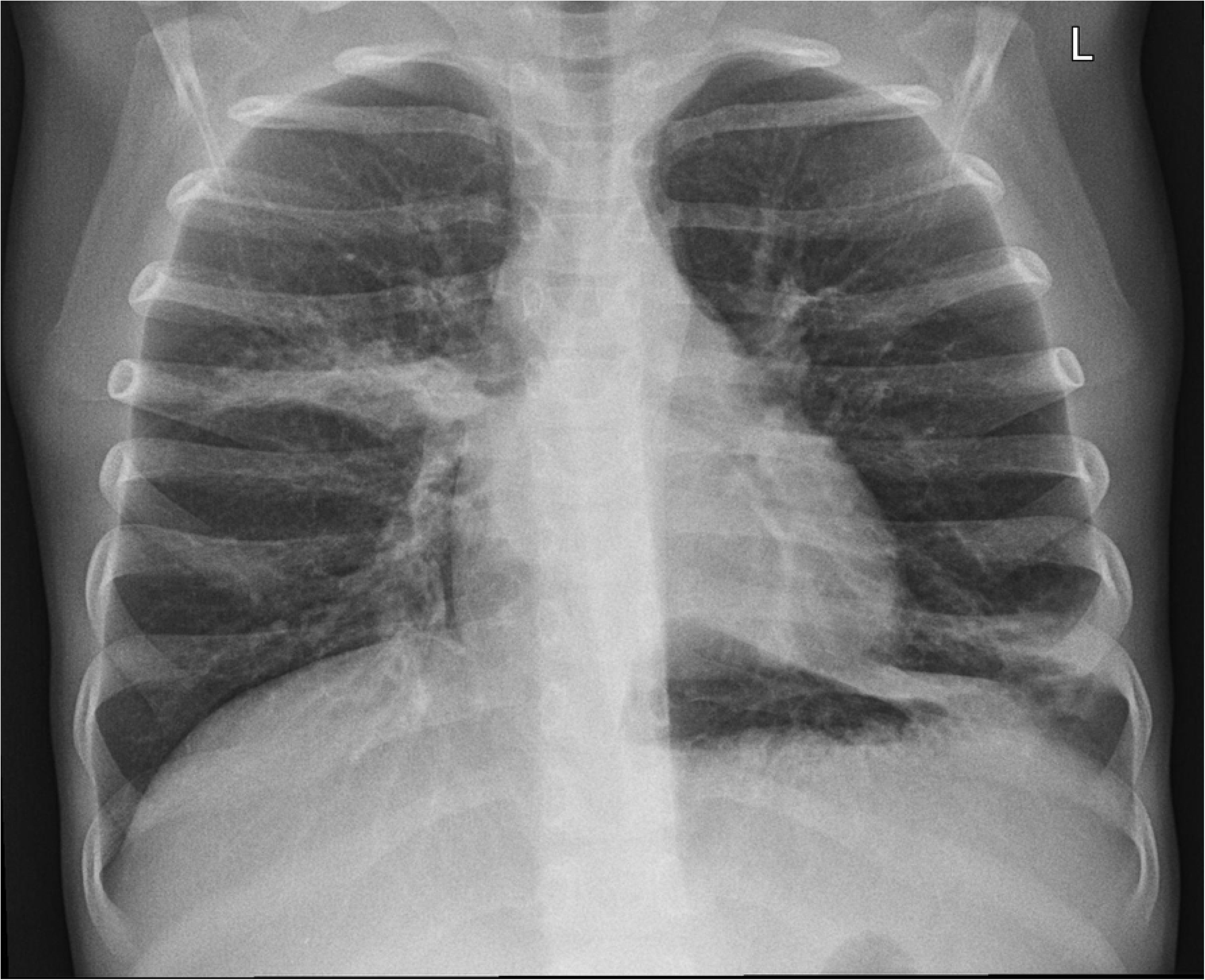
Linear opacities and regions of consolidations in the middle right and lower left field. Fine peribronchial infiltrates in the lower right and middle left field. Hyperinflation of both lungs. 15 points in modified Chrispin-Norman score.

**Fig 10.**
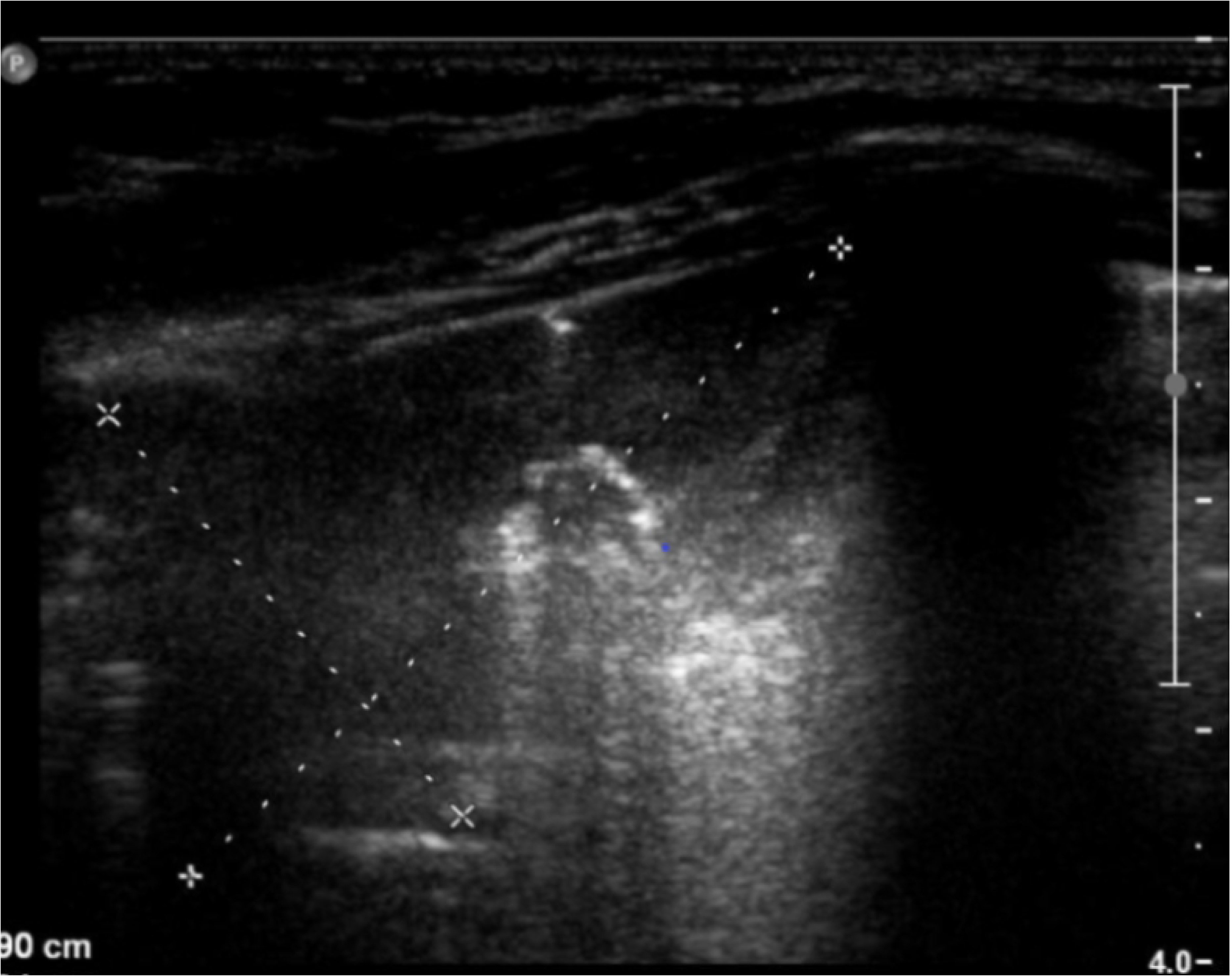
LUS image of the patient from Fig 7, linear probe. Large area of consolidation in the right upper lobe. 5 points in CF-USS.

**Fig 11.**
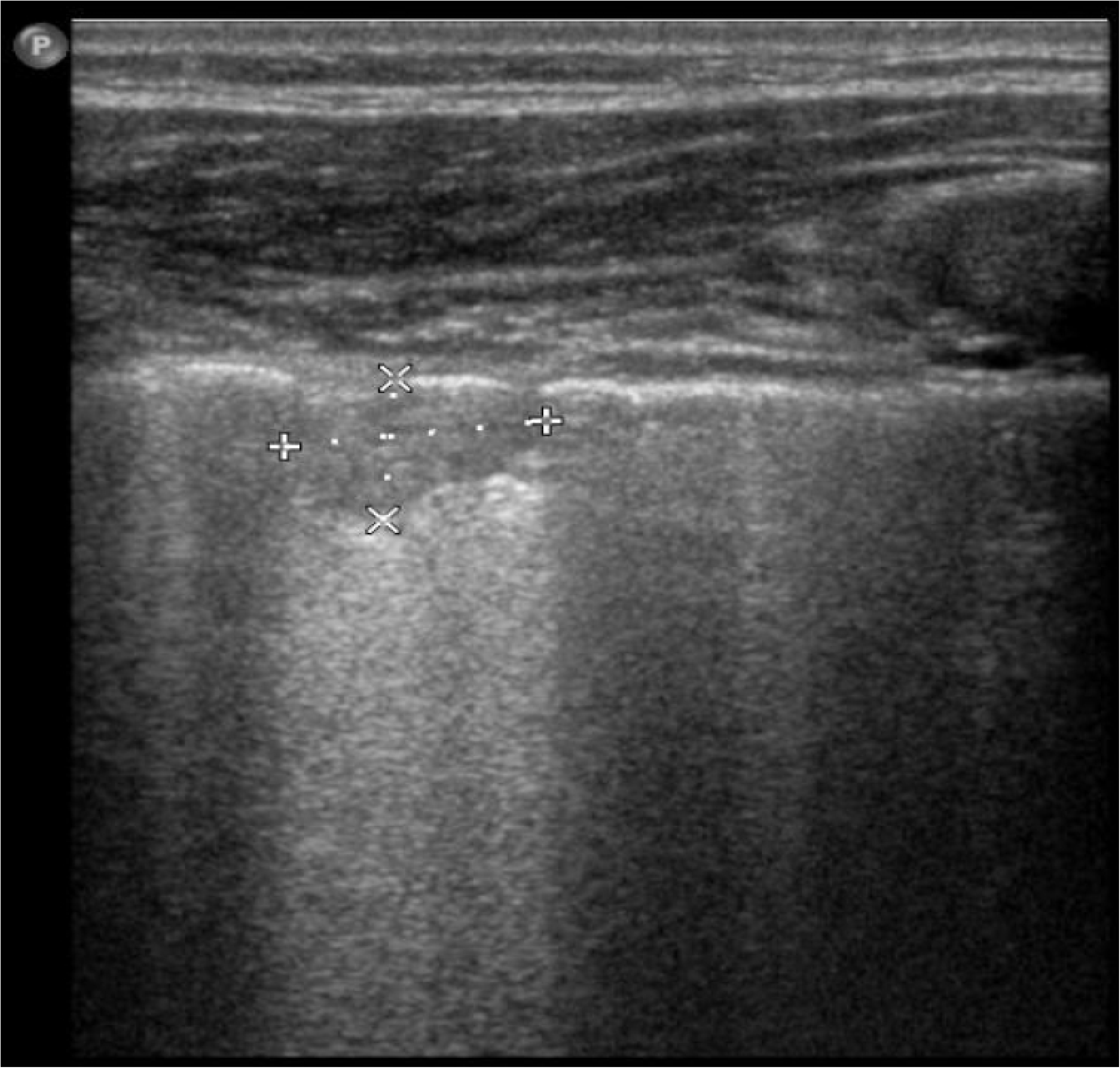
LUS image of the patient from Fig 7, linear probe. Fine subpleural consolidation.

## Discussion

Cystic fibrosis is a life-shortening genetic disorder, involving respiratory system and requiring chronic therapy. In the course of the disease patients suffer recurrent exacerbations, that affect patients quality of life and survival. Radiology plays a significant role in patients’ follow-up, enabling monitoring of the disease, response to treatment as well as the diagnosis of exacerbations [41]. Unfortunately conventional x-rays, especially numerous, repeated in the course of disease add up to cumulative ionising radiation dose. Diminishing radiation exposure, by looking for alternative diagnostic modalities, should be considered as one of the goals of contemporary medicine.

Chest ultrasound has not been routinely used in the monitoring of lung disease in paediatric patients with CF, despite its lack of radiation, availability and safety. Considering this we wanted to compare diagnostic value of LUS with conventional chest x-rays assessed according to the modified Chrispin – Norman score. There are no other studies comparing modified Chrispin – Norman score with chest ultrasound in CF paediatric patients. Furthermore, we developed our own ultrasound scoring system: CF-USS to make the comparison more feasible

For the evaluation of chest x-ray we chose modified Chrispin – Norman score as it uses only antero – posterior projections for the evaluation of the hyperinflation of the chest based on the shape of the thorax, diaphragm location and lung hyperlucency resulting from air-trapping [9]. That stays in agreement with the Benden et al. data and allows for the diminished radiation dose while avoiding the lateral projection [9]. Terheggen-Lagro and colleagues in their study compared six different clinical and radiological scoring systems (Schwachman – Kulczycki score, Chrispin-Norman score, modified Chrispin-Norman score, Brasfield score, Wisconsin score and Northern score) and demonstrated their clinical utility in different clinical settings [14]. Authors proved, that radiographic scoring systems in the CF patients, especially modified Chrispin-Norman score are characterized by low interobserver variability and correlate with pulmonary function tests results as well as clinical features.

The aim of the study was to compare the results of x-ray scoring system with chest ultrasound scoring system. Furthermore we constructed a novel chest ultrasound score for the evaluation of CF paediatric patients (CF-USS). The score has been developed based on the experience of Caiulo and colleagues, who used LUS in patients with bronchiolitis, the pathology, that among others is also present in the CF patients [5,15,40,42].

LUS was performed at the same time as the chest x-ray. In nine patients LUS were performed twice on two different occasions. The most commonly seen pathological features were B-line artefacts of different number and intensity. B-line artefacts might be seen in a normal lung and are not considered pathological as long as their number does not exceed 2 in a single transverse scan with a convex probe and 6 in a single longitudinal scan with a high resolution linear probe [32].

Clinical relevance of B-line artefacts is quite wide and has recently been covered in an excellent review by Dietrich and colleagues [43]. The authors believe, that B-line artefacts can be caused by multiple factors, and be present in lung oedema, heart failure, lung interstitial diseases, infections, acute respiratory distress syndrome (ARDS) or lung injury. B-line artefacts are the sign of increased lung density due to the loss of the lung tissue aeration. Chiesa and colleagues found B-line artefacts in 37% of elderly studied as compared to 10% of healthy young adults [44]. Correct B-line artefacts interpretation should account for the evaluation of other LUS signs and clinical data. In the pathological conditions B-line artefacts may be useful for the monitoring of treatment. The influence of technical factors on the appearance of B-line artefacts still remains to be elucidated [43].

In our LUS scoring system (CF-USS) B-line artefacts are divided into focal (few, some and many) and coalescent (absent, fused and “lung rockets”). The scoring system reflects intensity and variability of the lung pathology, known as B-line artefacts. Despite statistically significant correlation between the two studied scores in our material, we believe that true clinical significance of B-line artefacts, in a single LUS examination without clinical data has important limitations. In the children with CF B-line artefacts should be evaluated in the context of disease progression, documenting their numbers and localisation [45].

Another pathology seen in LUS are subpleural consolidations. Very fine 3 to 4 mm in diameter subpleural consolidations may be present in healthy children in the first few years of life [20,46]. In children with CF however they have important clinical implications. Dense mucus is blocking the airways, including bronchiole, leading to focal atelectasis and hyperinflation, and resulting in recurrent infections. Peripheral mucus plugs are causing small foci of inflammation, that might progress into disease exacerbations [10]. There are several studies illustrating the fact, that structural changes seen in radiologic examinations may be seen ahead of pulmonary function deterioration [47]. In CF-USS subpleural consolidations were classified as: absent, few and multiple or extensive. In 75% of the studied patients subpleural consolidations were seen in LUS and in 17% the changes were multiple or wider than 10 mm. None of the changes smaller than 10 millimetres were seen in conventional radiograms. In our opinion subpleural consolidations, similarly to B-line artefacts cannot be evaluated without clinical data. Brody and colleagues reported, that in stable CF patients subpleural consolidations should be monitored, as they may lead to clinical deterioration and decline in pulmonary function test results [47]. In patients with bronchopulmonary disease exacerbations diagnosing and monitoring of subpleural consolidations with LUS may limit the number of x-rays performed and in consequence radiation exposure.

CF-USS also comprised the evaluation of pleural fluid which seems reasonable to perform in patients suspected of pleural complications regardless of conventional x-ray results. In the studied group in the 23% of patients we have documented the presence of fluid in the pleural space, majority of them having just the small amount in the costo – phrenical angle. Pleural irregularities were seen in only one of the studied patients. Caiulo and colleagues reported in their bronchiolitis study pleural irregularities in 25% of patients that have been disappearing in the course of follow up [40]. The reason for the disparity of our results might be the fact, that bronchiolitis is not present an all patients with cystic fibrosis.

In 9 patients we repeated LUS and conventional x-rays in stable condition in the course of 2 years follow up. There were no statistically significant differences between either LUS or x-rays. Spearman rank correlation coefficient between x-ray and LUS was higher in the second series of studies.

We believe that LUS is an important diagnostic tool, accessory to conventional x-rays enabling monitoring of the disease process. We do acknowledge however the limitations of CF-USS scoring system. Our material is small and the study was conducted in a single CF centre – we do hope however, that the study will be continued in the future. The most important limitation remains inability of visualisation of consolidations separated from pleura as well as of airway pathologies (bronchiectasis, mucus plugs) that constitute the mainstay of lung pathology in CF pulmonary disease. Nevertheless, due to its safety, non-invasiveness and availability we hope that CF-USS will find its place in long term monitoring of the disease, response to treatment and the risk of exacerbations

## Conclusions

1. LUS should be an accessory radiographic examination in the scheduled follow-up visits in cystic fibrosis paediatric patients, and CF-USS scoring system may provide clinicians with valuable informations concerning the disease progression and exacerbations’ risk.
2. CF-USS results correlate with conventional x-ray modified Chrispin – Norman score.
3. LUS constitute an invaluable tool for the diagnosis of subpleural consolidations
4. LUS limitations remain inability to visualise consolidations separated from the pleura and larger airways. Numerous clinical conditions in which B-line artefacts can be present additionally makes it difficult to recommend LUS as the sole diagnostic modality in cystic fibrosis patients.

Authors have no conflict of interest

